# FOXM1 regulates platelet-induced anoikis resistance in pancreatic cancer cells

**DOI:** 10.1101/2024.10.31.620459

**Authors:** Alissa Ernesti, Beate Heydel, Juliane Blümke, Tony Gutschner, Monika Hämmerle

## Abstract

Anoikis resistance is a prerequisite for circulating tumor cells (CTC) to survive in the blood stream. Platelets can interact with these CTCs and protect them from cytokine and immune cell-mediated cell death. Whether platelets can regulate anoikis resistance by controlling tumor cell intrinsic gene expression changes has not been studied in pancreatic cancer cells in detail before. Here, we identified FOXM1 as a differentially regulated gene between attached and detached cells whose expression was controlled by platelets. Manipulating FOXM1 expression using FOXM1 knockdown and overexpression or by inhibiting its function using a small molecular inhibitor highlighted the role of FOXM1 in controlling platelet-mediated anoikis resistance in pancreatic cancer cells. Hence, targeting FOXM1 might be a novel therapeutic strategy in pancreatic cancer patients, especially in those with thrombocytosis.

## Introduction

Pancreatic cancer is a deadly disease with a poor overall survival rate. As of 2022, pancreatic cancer is the sixth leading cause of cancer-related death worldwide with highest incidence rates in Europe, North America, and Australia (Bray et al., 2024). Most patients die due to late diagnosis, already having local or distant metastases (Kleeff et al., 2016). Metastasis is a highly controlled process involving detachment of cancer cells from the primary tumor, intravasation into blood and lymphatic vessels, extravasation into distant organs and ultimately the ability to form a secondary mass (Gerstberger et al., 2023). For successful formation of distant metastases, survival of circulating tumor cells (CTCs) in the blood is crucial. During their short transit time, CTCs encounter several challenges that impact their survival including shear stress and contact with immune cells including natural killer cells. It has been suggested that survival of CTCs can be greatly enhanced by their interaction with platelets forming a platelet coat on the CTC surface and physically shielding cancer cells from immune and physical stressors (Labelle and Hynes, 2012). These microclusters of platelets and tumor cells have a high metastatic potential, as various studies have shown that platelets are able to promote metastasis formation in cancer (Haemmerle et al., 2017; Labelle et al., 2011). However, whether platelets are able to enhance survival of pancreatic cancer cells under low attachment conditions that would contribute to metastasis formation has not been studied in detail. Here, we sought to determine the underlying mechanisms that are relevant for the survival of pancreatic cancer cells after detachment and evaluated the effects of platelets as mediators of anoikis resistance. Based on these analyses, we identified FOXM1, regulated by Akt, as a crucial regulator of survival of detached cells and of platelet-mediated gene expression changes and showed that FOXM1 is critical for platelet-induced anoikis resistance in vitro.

## Material and Methods

### Cell culture

Cell lines used in this study were obtained from ATCC or DSMZ. IMIM-PC1 cells were a gift from Prof. Michl (Internal Medicine I, University Hospital Halle). Cell lines were routinely tested for mycoplasma contamination using mycoplasma-specific PCR. Human pancreatic cancer cell lines BxPC-3, SU.86.86 and AsPC-1 were cultured in RPMI-1640 supplemented with 10% fetal bovine serum (FBS) and 1% penicillin-streptomycin. Human pancreatic cancer cell lines PA-TU-8988T, PA-TU-8988S, CAPAN-1, PANC1, IMIM-PC1 and MIA PaCa-2 were cultured in DMEM supplemented with 10% fetal bovine serum (FBS) and 1% penicillin-streptomycin. Cells were maintained at 37°C in a humidified incubator infused with 20% O_2_ and 5% CO_2_.

### siRNA transfection

For experiments involving small interfering RNA (siRNA) transfections, control siRNAs and two individual FOXM1 siRNAs were transfected (40nM f.c. siRNA) using RNAiMAX (Thermo Fisher Scientific) 24 hours before collecting cells and seeding them into low-attachment plates. Thereafter, cells were co-incubated with platelets and collected 48 hours later for RNA expression analysis and propidium iodide (PI) staining and flow cytometry analysis. Successful knockdown after transfection and following 48 hours of low attachment cultures was checked using qRT-PCR. The following siRNA have been used in this study: AllStars Neg. Control siRNA (Qiagen), FOXM1 siRNA 1: AUAUUCACAGCAUCAUCAC, FOXM1 siRNA 2: GGACCACUUUCCCUACUUU.

### Generation of FOXM1 overexpressing BxPC-3 cells

FOXM1 overexpression plasmid was generated by cloning the FOXM1 coding sequence into a modified pHAGE vector (modified pHAGE-SMAD4 vector; #116791 from Addgene; kind gift from Gordon Mills & Kenneth Scott) (Ng et al., 2018). The SMAD4 insert was replaced by a multiple cloning site containing BsrGI, NcoI, NheI, BmtI, NotI, PspXI, XbaI, BstBI and MluI. FOXM1 coding sequence including a Flag-tag was used from a Flag-FOXM1 plasmid (#153136 from Addgene; kind gift from Stefan Koch) (Moparthi and Koch, 2020) and cut out using NcoI and XbaI restriction enzymes. Lentivirus particles were produced using HEK293T, cultured in DMEM, with packaging and VSV-G envelope expressing plasmids (#12259 and #12260 from Addgene; kind gift from Didier Trono). HEK293T supernatants were harvested 48 hours after transfection and BxPC-3 cells were transduced immediately using fresh virus particles. 24 hours after transduction, BxPC-3 cells were selected using 1 µg/mL of Puromycin (cat. no. A1113803, Thermo Fisher Scientific). Overexpression was checked on RNA and protein level using qRT-PCR and western blotting.

### Isolation of murine platelets

Platelets were isolated as previously described (Haemmerle et al., 2017). Briefly, whole blood was drawn from the inferior vena cava of anaesthetized nude or C57BL/6 mice into a 1 ml syringe that has been pre-loaded with 100 µl of acid citrate dextrose solution B (GT Biosciences). Afterwards, the blood was gently mixed with 500 µl of tyrodes buffer. Blood was centrifuged at 120 g for 6 minutes at room temperature. The platelet-rich plasma fraction was passed through a filtration column of sepharose CL-2B beads (GE Healthcare) loaded into a siliconized glass column with a 10 µm nylon net filter (Millipore). Cloudy eluents containing platelets were collected in buffer I using a 15 ml falcon tube. The number of platelets was counted with a hemocytometer and immediately used for subsequent experiments. All animal experiments were conducted in accordance with the file reference MLU 2-1584 (Landesverwaltungsamt Saxony-Anhalt).

### Low attachment/anoikis assay in vitro

On day 1, 500,000 cells were seeded in an ultra-low attachment 6-well plate (cat. no. 3471, Corning). Immediately thereafter, platelets were isolated from the blood of nude or C57BL/6 mice and 100 × 10^6^ platelets were added to cancer cells. 48 hours after platelet addition, cells were lysed for RNA or protein analysis. 48h or 72h after platelet addition the number of PI - positive cells was evaluated by a MACSQuant® Analyzer 10 (Miltenyi Biotec). In indicated experiments, 5 µM of FOXM1 inhibitor FDI-6 (cat. no. S9689, Selleck Chemicals) or 1 µM AKT inhibitor MK-2206 (cat. no. S1078, Selleck Chemicals) was added at time of cell seeding.

### RNA isolation, cDNA synthesis, and real-time PCR analysis

Cells were harvested using 1 mL TRIZOL as previously described (Dorn et al., 2020). Isolated RNA was resuspended in 20-30 µL ultrapure water. For cDNA synthesis up to 1 µg of total RNA was reverse transcribed using random hexamer primers and M-MLV Reverse Transcriptase (Promega). Gene expression was measured using the primaQUANT CYBR qPCR mastermix (Steinbrenner Laborsysteme GmbH) with a Light Cycler® 480 II (Roche). Sequence information for qRT-PCR primers are listed in Table 1.

**Table 1:**
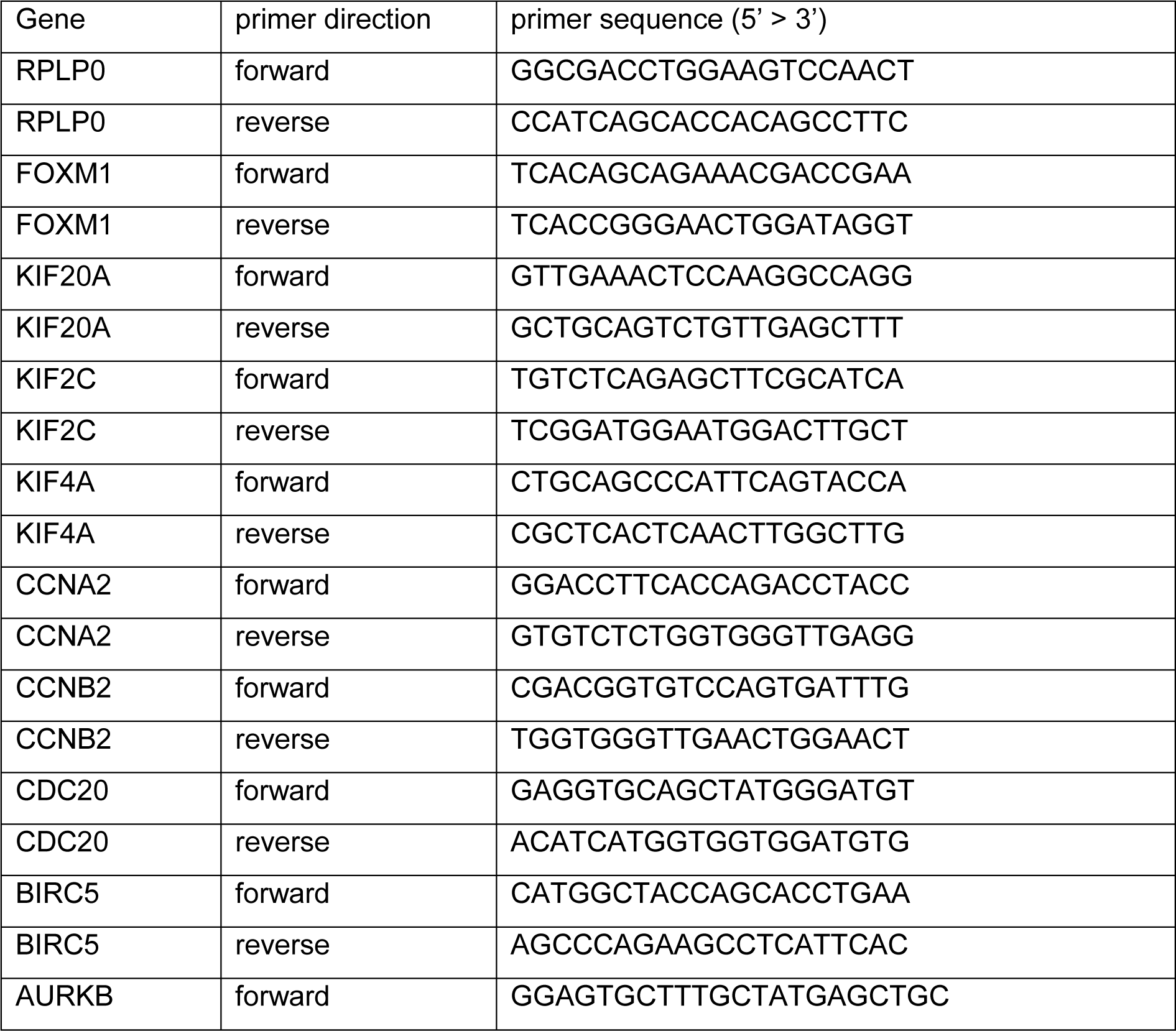
List of qRT-PCR primers used in this study.

### mRNA-sequencing

RNA was isolated as described above from BxPC-3 and SU.86.86 cells under low attachment or attached condition with or without platelet co-incubation for 48h. mRNA-sequencing was performed with 2 µg of total RNA (n = 3) for each sample. Library preparation and sequencing was performed by Azenta (Leipzig, Germany). Library preparation was based on polyA-tail selection and sequencing was performed on an Illumina NovaSeq platform resulting in approximately 20 million reads per sample. Trim Galore! v0.4.3.1 was used for quality check (80% bases Q ≥ 30). Mapping to the human genome hg38 followed by differential gene expression analysis was performed using RNA STAR v2.6.0b-2 and edgeR v 3.24.1.

### Gene set enrichment analysis and protein-protein-interaction prediction

Differentially expressed genes were pre-ranked according to their log2 fold change and analyzed using the GSEA software (Mootha et al., 2003; Subramanian et al., 2005). Results with a FDR q-value of ≤ 0.05 were considered statistically significant. Furthermore, prediction of protein-protein interactions was performed using STRING (https://string-db.org) with a minimum required interaction score of 0.7 (Szklarczyk et al., 2023).

### Western blot analysis

Cells were harvested by lysing the cell pellet using RIPA buffer (50 mM Tris-HCl pH 8.0, 150 mM NaCl, 1% (v/v) IGEPAL CA-630, 0.5% (w/v) Na-deoxycholate, 0.1% (w/v) SDS), supplemented with 10% protease and phosphatase inhibitors (cat. no. A32957, Thermo Fisher Scientific, cat. no.11836153001, Roche). SDS-polyacrylamid gel electrophoresis (SDS-PAGE) was performed at 120V and proteins were transferred onto a nitrocellulose membrane using the wet blot method. Detection of protein expression was performed using Bio-Rad ChemiDoc MP (Bio-Rad Laboratories). Quantification of protein expression was done using ImageJ (Schneider et al., 2012). The following primary antibodies were used: FOXM1 (clone D12D5; #5436), PARP (clone 46D11; #9532), cleaved PARP (clone D64E10; #5625), phospho-Akt (Ser473, clone D9E; #4060), Akt (clone C67E7; #4691), all from Cell Signaling Technology, Danvers; GAPDH (clone GAPDH-71.1; #G8795) from Sigma-Aldrich Chemie GmbH; RPL7 (#A300–741A; Thermo Fisher Scientific, Waltham). The secondary antibodies (IRDye® 800CW Donkey anti-Rabbit; IRDye® 680RD Donkey anti-Mouse) were fluorescently labelled and purchased from LI-COR Biosciences GmbH.

### Immunofluorescence

Suspension cells were spun onto a glass microscope slide that has been pre-treated with poly-L-lysine (cat. no. P8920, Sigma-Aldrich Chemie GmbH) at 5000 rpm for 5 minutes. Slides were dried for another five minutes. Thereafter, cell were fixed in 4% paraformaldehyde for 30 min. followed by washing in PBS and incubation with blocking buffer (1X PBS / 1 % BSA / 0.3% Triton™ X-100) for 60 min. Incubation with FOXM1 primary antibody, diluted 1:100 in blocking buffer (clone D12D5; #5436; Cell Signaling Technology) was performed overnight at 4°C. On the next day, slides were washed with PBS and incubated for 1 h with a secondary α-rabbit AlexaFluor® 488 antibody (Jackson ImmunoResearch Laboratories), diluted 1:500 in blocking buffer. Actin cytoskeleton was visualized using Phalloidin staining (P1951; Sigma-Aldrich Chemie GmbH) and nuclei were stained with DAPI (1 µg/mL; Carl Roth). After a final wash, slides were mounted using Mowiol (Calbiochem). Pictures were taken using a Keyence BZ-X microscope.

### Statistical analysis

Statistical analysis was done using Excel and GraphPad Prism 8. Differences between groups were evaluated using two-tailed Student’s t-test or one-way analysis of variance (ANOVA), adjusting for multiple comparisons. Results are presented as the mean ± SEM. Survival analysis has been done using R2: Genomics Analysis and Visualisation Platform (https://r2.amc.nl). For all statistical analyses, p ≤ 0.05 was considered statistically significant.

## Results

### Platelets induce anoikis resistance in pancreatic cancer cells

Survival of cancer cells in the blood stream and resistance to cell death occurring after detachment from the extracellular matrix is essential for subsequent metastasis formation. To model non-adherent cultures and examine anoikis rates of human pancreatic cancer cells, we incubated human cell lines using ultra-low attachment cell culture plates. We decided to use 72 hours after cell seeding as the major endpoint for measuring anoikis levels *in vitro* as published before (Haemmerle et al., 2017). In all cell lines, we measured a significant number of dead cells detected by PI staining (**Figure 1A-C, Supplementary Figure 1A-E, upper panels**). Interestingly, individual differences in anoikis rates between cell lines were detected. While PA-TU-8988T (**Supplementary Figure 1D**) and AsPC-1 cells (**Supplementary Figure 1E**), both cell lines derived from either liver metastasis or ascites, were highly resistant to anoikis with only 20% PI-positive cells after 72 h, BxPC-3, Su86.86 and IMIM-PC1 cells (**Figure 1A-C**) were very sensitive to the low attachment culturing condition with apoptosis rates between 75% and 90%. In addition to the flow cytometry results, increased apoptosis rates were confirmed using western blot analyses of the respective cell lines under attached and low attachment conditions. Specifically, high signals for cleaved PARP were detected for BxPC-3, Su.86.86 and IMIM-PC1 (**Figure 1A-C, lower panels**), while only minor differences in low attachment compared to attachment cultures were observed in PA-TU-8988T and AsPC-1 cells (**Supplementary Figure 1D-E, lower panels**). As platelets have been shown to facilitate survival of ovarian and colorectal cancer cells in the blood stream and thereby support metastasis formation (Haemmerle et al., 2018; Haemmerle et al., 2017), we examined whether platelets could improve survival of pancreatic cancer cells under low attachment conditions. Therefore, we co-cultured cells that were initially sensitive to the anoikis condition with 100 x 10^6^ platelets and measured the percentage of PI-positive cells after 72 hours. Interestingly, co-incubation of BxPC-3, Su.86.86 and IMIM-PC1 cells with platelets significantly decreased the number of dead detached cancer cells in all the tested cell lines and increased survival rates up to 60% (**Figure 1D-F, upper panels**). In addition, cleaved PARP levels decreased markedly after tumor cell – platelet co-incubation (**Figure 1D-F, lower panels**).

**Figure 1:**
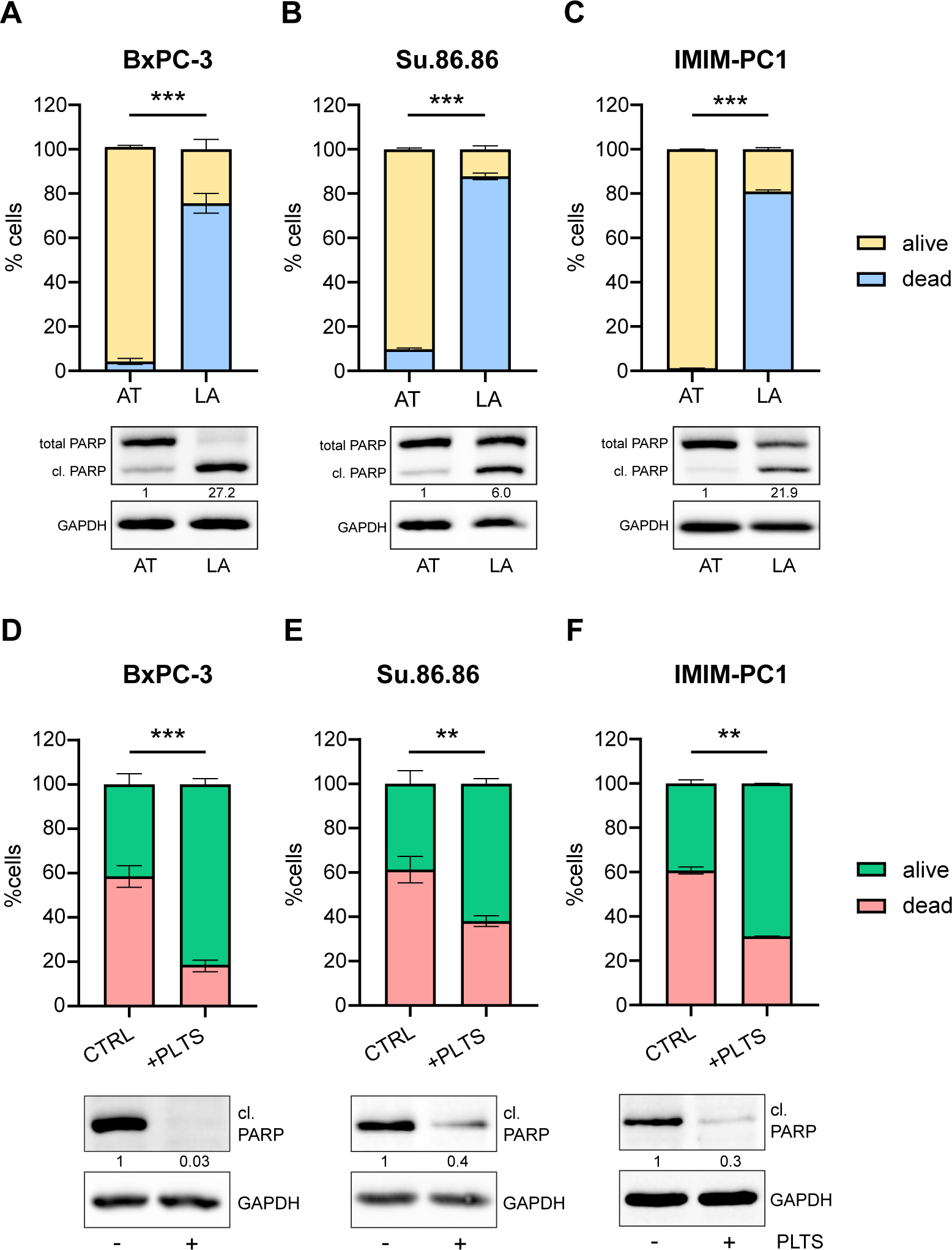
Anoikis rates change under low attachment conditions and platelet co-incubation. Pancreatic cancer cells were cultured under attached (AT) and low-attachment (LA) conditions for 72 hours and the % of dead (PI-positive) and living (PI-negative) BxPC-3 **(A)**, Su.86.86 **(B)** and IMIM-PC1 **(C)** cells were measured using flow cytometry (upper panel). In addition, increased apoptosis rates were confirmed using protein analysis and quantification of cleaved versus full length PARP (lower panel). GAPDH was used as a loading control. **(D-F)** Respective cell lines were co-incubated with 100 x 10^6^ platelets (PLTS) and flow cytometry analysis and western blotting was done as described above. Bars and error bars represent mean values and the corresponding SEMs (n=3; **p < 0.01, ***p < 0.001).

### Gene expression signatures upregulated in attached cells and by platelets

Next, we sought to analyze signaling pathways responsible for cancer cell survival under attached and low-attachment conditions and the effect of platelets on gene expression program in detached cells. As BxPC-3 and Su.86.86 cells were highly sensitive to the anoikis condition and significantly responded to platelet co-incubation, we performed RNA expression analysis of these cell lines and first compared RNA signatures under attached versus low-attachment conditions. We found 265 genes and 184 genes differentially regulated between adherent and non-adherent BxPC-3 and Su.86.86 cells, respectively. Overlapping gene expression changes of both cell lines revealed a common regulation of 109 genes (**Figure 2A**). To identify gene expression changes and molecular pathways that were regulated by platelets in cancer cells, we were specifically interested in genes that were downregulated in non-adherent cells compared to attached conditions and again upregulated in cells that interacted with platelets. Overlapping these gene lists identified a set of 24 genes regulated in both cell lines and by platelet co-incubation (**Figure 2A, B, Supplementary Figure 2A**). Gene Set Enrichment Analysis (GSEA) showed that pathways related to cell cycle and E2F1-signaling were the most upregulated pathways in cells co-incubated with platelets, both in BxPC-3 (**Figure 2C**) and Su.86.86 cells (**Supplementary Figure 2B**). Intriguingly, string analysis of the 24 overlapping genes highlighted one prominent cluster of genes displaying strong protein-protein interactions (**Figure 2D**).

**Figure 2:**
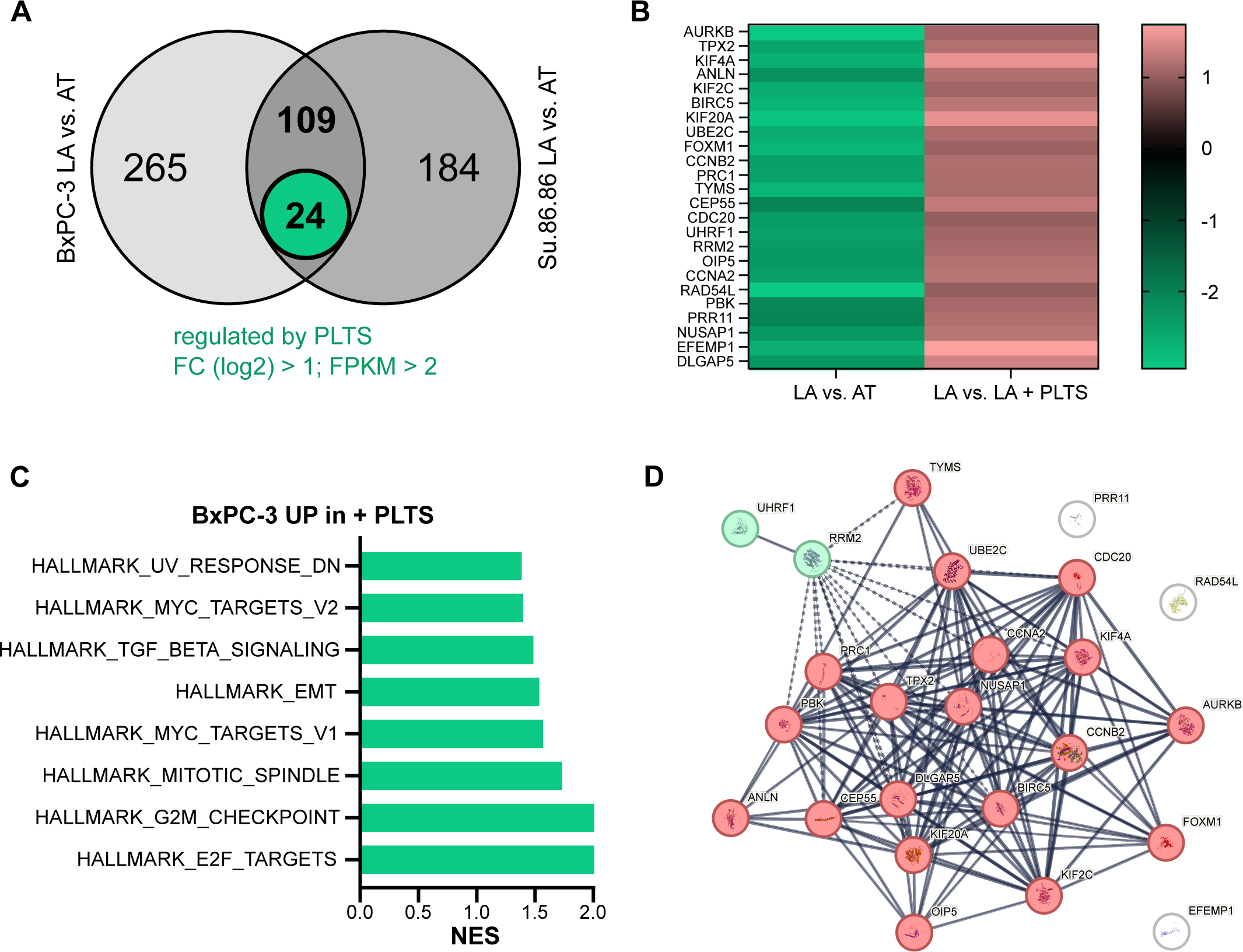
Platelets induce a FOXM1-specific gene signature in cancer cells. **(A)** RNA sequencing analysis of cells under low attachment (LA) versus attached (AT) conditions revealed 265 and 184 differentially regulated genes in BxPC-3 and Su.86.86 cells, respectively. Overlapping gene lists with sequencing results of cells co-incubated with platelets revealed a common regulation of 24 genes in both cell lines. Fold change (FC; log2) cut-off was 1; FPKM cut-off was 2; FDR ≤ 0.05 (n=3). **(B)** Enriched pathways of upregulated genes in platelet co-incubated BxPC-3 cells using Gene Set Enrichment Analysis (GSEA, www.broadinstitute.org/gsea). **(C)** Heat map showing 24 differentially regulated genes in BxPC-3 cells. **(D)** STRING analysis showed protein-protein interactions between regulated genes based on experiments, databases, co-expressions and co-occurrence (string-db.org).

### FOXM1 expression is downregulated in non-adherent cells and associated with anoikis

One important transcription factor downregulated in non-adherent cells, but upregulated by platelets was forkhead box transcription factor M1 (FOXM1). FOXM1 is a regulator of cell cycle associated genes and essential for DNA replication and mitosis (Laoukili et al., 2005; Wang et al., 2005). In cancer, FOXM1 is claimed to be a master regulator of tumor metastasis (Raychaudhuri and Park, 2011), is highly expressed in several cancers and associated with poor patient survival (Liao et al., 2018). Recently, it has been shown that metastatic spread of ovarian cancer in the peritoneal cavity and ovarian cancer stemness is dependent on FOXM1 (Battistini et al., 2024; Parashar et al., 2020). In addition, expression of numerous other genes downregulated by detachment and upregulated by platelet co-incubation including KIF2C (Zhao et al., 2014), KIF20A (Fang et al., 2023), KIF4A (Hu et al., 2019), CCNA2 and CCNB2 (Vasudevan et al., 2018), CDC20 (Xie et al., 2015) and BIRC5 (Wang et al., 2005), were previously shown to be controlled by FOXM1. Re-analyzing PDAC datasets from Bailey et al. (Bailey et al., 2016) revealed that the most aggressive, namely squamous subtype of pancreatic cancer, showed the highest FOXM1 expression (**Supplementary Figure 3A**) and that FOXM1 expression was associated with worse overall survival (**Supplementary Figure 3B**). In order to show that FOXM1 RNA and protein expression was regulated by cellular detachment, qRT-PCR as well as western blot analyses were performed. In all pancreatic cancer cell lines tested, except PA-TU-8988T, FOXM1 was significantly downregulated under low attachment conditions on the RNA level (**Figure 3A**). Similar results were obtained for FOXM1 protein levels (**Figure 3B**). In addition, in BxPC3 (**Figure 3C**) and Su.86.86 cells (**Supplementary Figure 4A**) qRT-PCR analysis confirmed RNA sequencing results of differentially expressed genes between attached and low-attachment conditions. Genes were also regulated in IMIM-PC1 cells that were highly sensitive to the low attachment conditions (**Supplementary Figure 4B**), whereas in PA-TU-8988T cells, which were resistant to detachment and which did not show deregulation of FOXM1 mRNA levels, expression of potential FOXM1 target genes was not affected (**Supplementary Figure 4C**). Lastly, RNA expression changes of FOXM1 depicted as ΔΔCt between adherent and non-adherent significantly correlated with percentage of PI-positive cells after 72 hours of low attachment (**Figure 3D**), suggesting that FOXM1 controls anoikis-mediated cell death under low attachment conditions.

**Figure 3:**
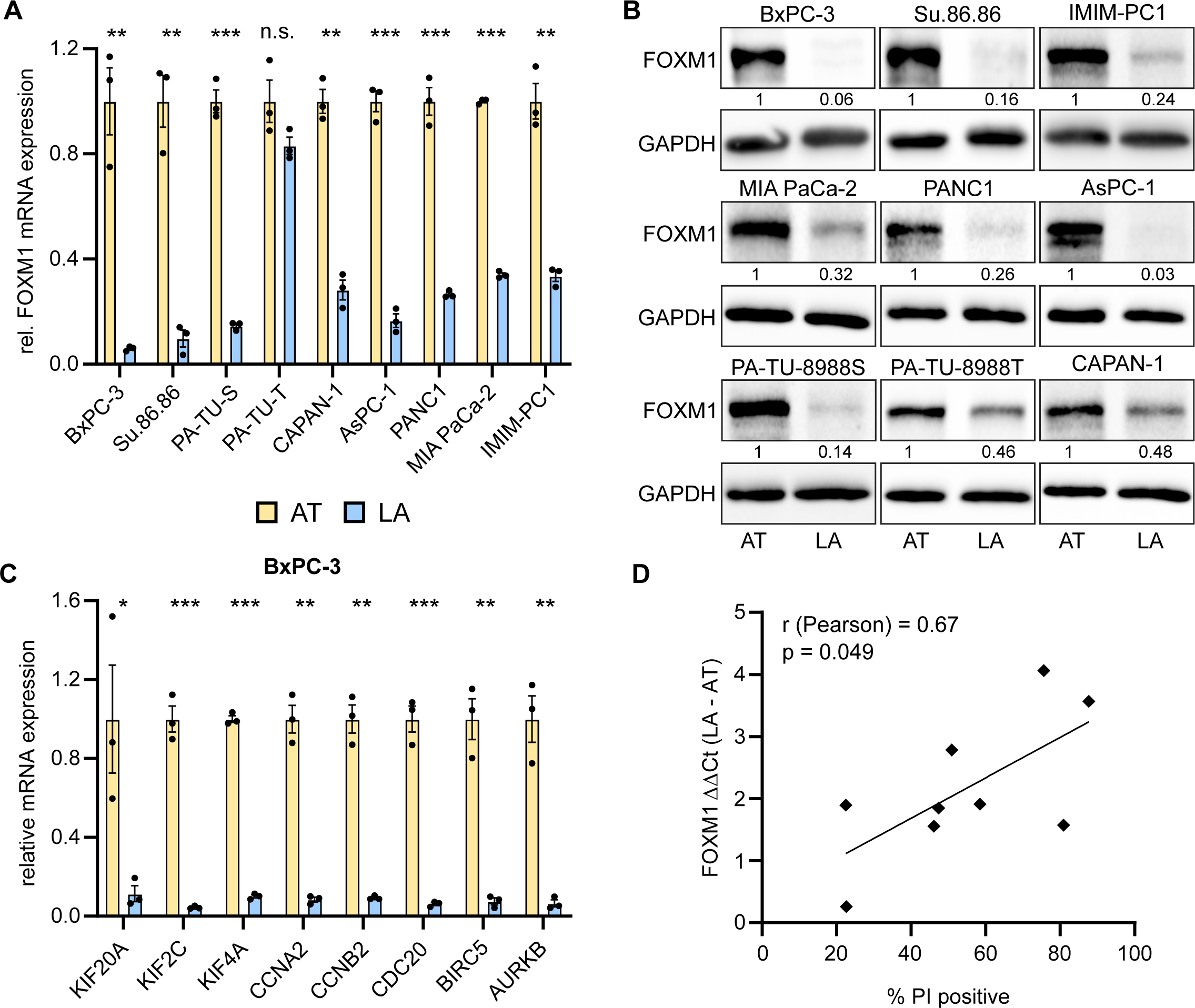
FOXM1 expression and associated gene signature is upregulated in attached cells. **(A)** FOXM1 mRNA expression is significantly higher in attached (AT) versus low-attachment (LA) cultures in all tested cell lines except PA-TU-8988T cells. RPLP0 was used as a reference gene. **(B)** Protein analysis and quantification of FOXM1 protein of attached and low-attachment cell cultures. GAPDH was used as a loading control. **(C)** Validation of RNA sequencing results of genes known to be regulated by FOXM1 in BxPC-3 cells using qRT-PCR. RPLP0 was used as a reference gene. **(D)** Expression change of FOXM1 mRNA (RPLP0 normalized) was correlated to % of PI positive (=dead) cells under low attachment. Bars and error bars represent mean values and the corresponding SEMs (n=3; *p<0.05, **p < 0.01, ***p < 0.001, n.s. non-significant).

### FOXM1 expression signature is induced by platelets and required for platelet-induced anoikis resistance

In order to validate RNA sequencing results, we measured FOXM1 expression on RNA and protein level in BxPC-3, Su.86.86 and IMIM-PC1 cells that were previously co-incubated with platelets. qRT-PCR and western blot analyses confirmed a strong upregulation of FOXM1 mRNA and protein by platelets in cancer cells after 48 hours co-culture (**Figure 4A, B**). In addition, we measured RNA expression changes of potential FOXM1 target genes in BxPC-3 (**Figure 4C**), Su.86.86 (**Figure 4D**) and IMIM-PC1 cells (**Supplementary Figure 4D**) that were co-incubated with platelets for 48 hours. Indeed, platelet co-culture led to a consistent and mostly significant upregulation of FOXM1 target genes in all three cell lines. In order to confirm whether FOXM1 is responsible for anoikis resistance induced by platelets, BxPC-3 cells were co-incubated with platelets with and without FDI-6. FDI-6 is a highly specific FOXM1 inhibitor that was shown to inhibit the FOXM1 transcriptional program by displacing FOXM1 from its genomic targets (Gormally et al., 2014). Whereas FDI-6 did not have a significant effect on the anoikis rate itself (**Supplementary Figure 5A**), it significantly reduced platelet-mediated anoikis resistance in BxPC-3 cells (**Figure 5A**). While platelet co-incubation reduced percentage of PI-positive cells by up to 60%, this was strongly inhibited by FDI-6 showing only a reduction between 5 % and 20%. Importantly, FDI-6 co-incubation only slightly reduced FOXM1 protein expression (**Figure 5B**). However, immunofluorescence analyses indicated that nuclear expression of FOXM1 was induced by platelets and inhibited by the addition of FDI-6 (**Figure 5C, Supplementary Figure 5B**). In order to see whether additional genes found to be upregulated by platelets (see Figure 2C) were downregulated by chemical FOXM1 inhibition, qRT-PCR analyses were performed on these samples. Indeed, FDI-6 treatment of detached BxPC-3 cells reduced expression of genes related to cell cycle regulation that were induced by platelet-cancer cell interaction (**Figure 5D**). In order to validate FOXM1 involvement in platelet-induced anoikis resistance, we knocked down FOXM1 expression using two individual siRNAs (**Supplementary Figure 5C**) before low attachment cultures. Again, platelet co-incubation reduced the number of dead BxPC-3 cells by more than 50%.

**Figure 4:**
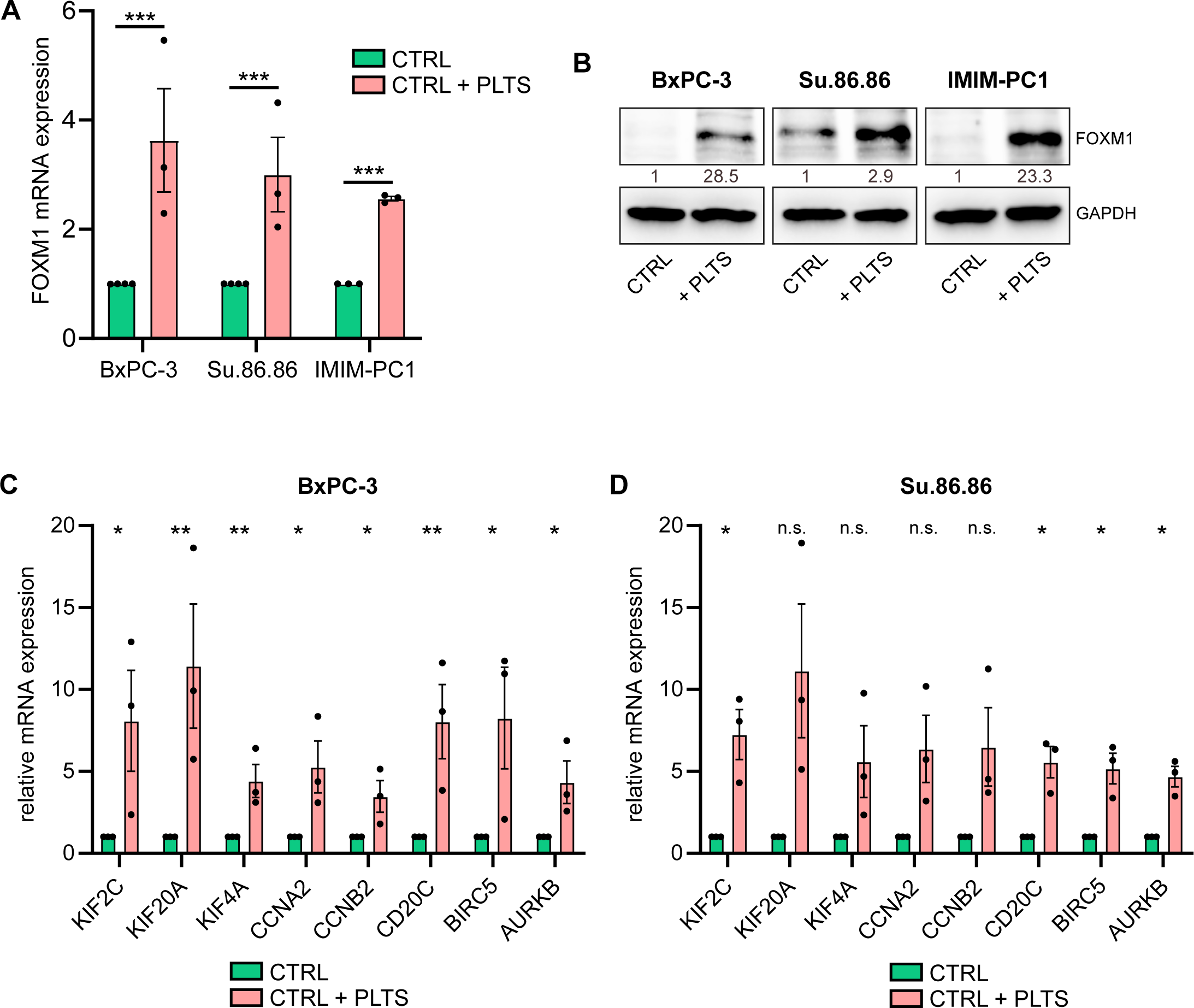
FOXM1 and FOXM1-associated gene signature is regulated by platelet co-incubation. qRT-PCR analysis **(A)** and western blotting **(B)** show increased FOXM1 mRNA and protein expression of FOXM1 after platelet co-incubation in BxPC-3, Su.86.86 and IMIM-PC1 pancreatic cancer cell lines. RPLP0 was used as a reference gene for qRT-PCR. GAPDH was used as a loading control for western blotting. **(C, D)** mRNA expression of FOXM1-regulated genes in BxPC-3 (C) and Su.86.86 (D) cells in low attachment cells co-incubated with platelets. Bars and error bars represent mean values and the corresponding SEMs (n=3; *p<0.05, **p < 0.01, ***p < 0.001, n.s. non-significant).

**Figure 5:**
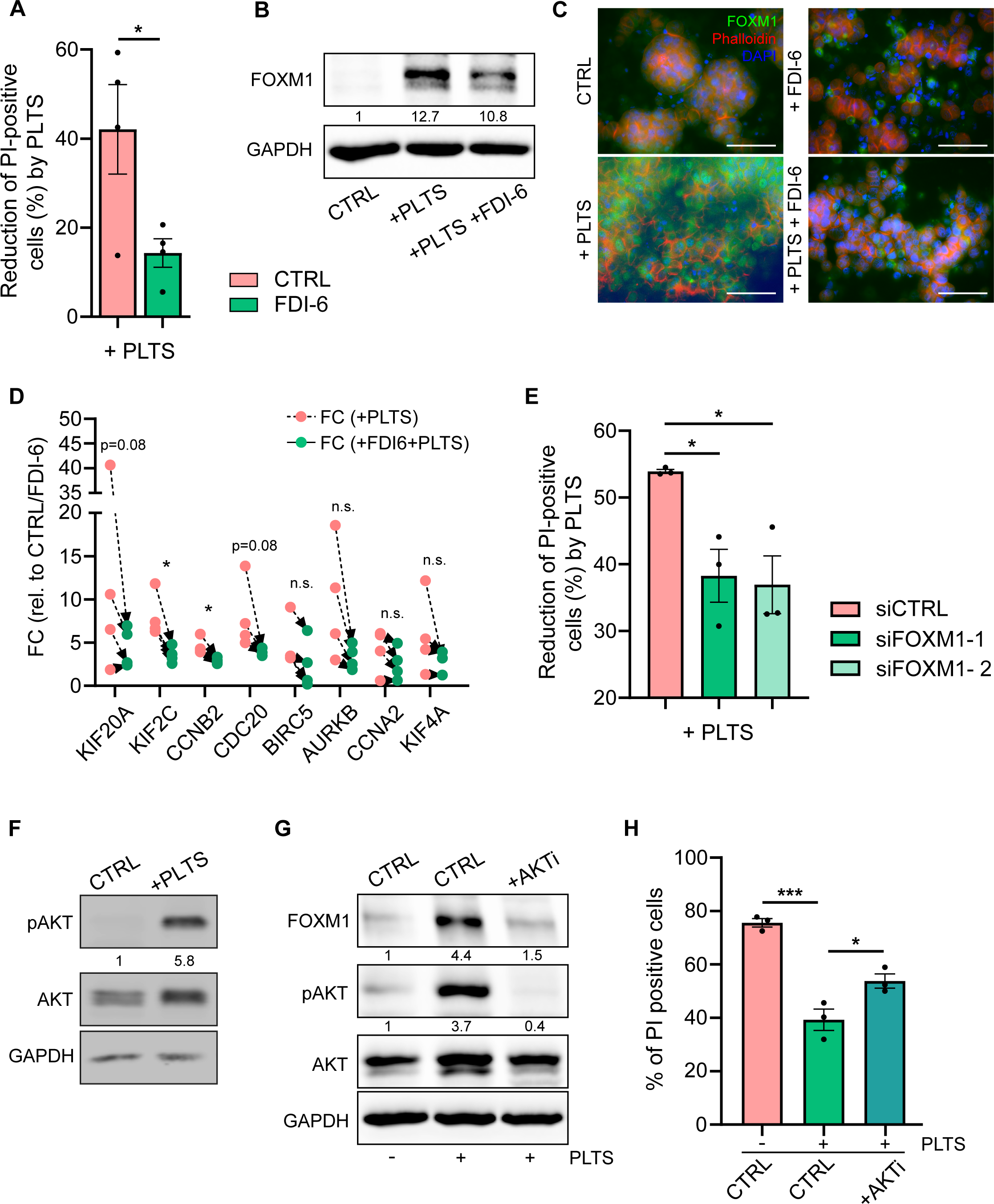
FOXM1 is necessary for platelet-mediated anoikis resistance. **(A)** Percentage (%) of reduction of PI-positive (dead) cells after platelet co-incubation of BxPC-3 cells with or without treatment with the FOXM1 small molecule inhibitor FDI-6. **(B)** Protein analysis and quantification of FOXM1 protein after 48 hours platelet co-incubation with or without FDI-6. GAPDH was used as a loading control. **(C)** Immunofluorescence of FOXM1 protein after platelet co-incubation with or without FDI-6. Cell cytoskeleton was visualized using Phalloidin. Nuclear counterstain was done using DAPI. Scale bar 50µm. **(D)** Gene expression changes after platelet co-incubation with or without FDI-6 (related to untreated control or FDI-6 treatment), measured by qRT-PCR. RPLP0 was used as a reference gene. **(E)** Percentage (%) of reduction of PI-positive (dead) cells after platelet co-incubation of BxPC-3 cells transfected with two individual FOXM1 siRNAs. **(F)** Protein analysis and quantification of phosphorylated AKT (Ser473) versus total AKT in BxPC-3 cells after 2 hours of platelet co-incubation. GAPDH was used as a loading control. **(G)** Protein analysis and quantification of phosphorylated AKT (Ser473) versus total AKT and FOXM1 protein after 48 hours of platelet co-incubation with or without AKT inhibitor MK-2206. GAPDH was used as a loading control. **(H)** Percentage of PI positive (dead) cells after 72 hours low attachment condition with or without platelet co-incubation or with AKT inhibitor MK-2206. Bars and error bars represent mean values and the corresponding SEMs (n=3; *p<0.05, **p < 0.01, ***p < 0.001, n.s. non-significant).

However, this effect was diminished by FOXM1 knockdown using both siRNAs (**Figure 5E**). Likewise, stable overexpression of FOXM1 in BxPC-3 (**Supplementary Figure 5D**) reduced PI-positive cells under low attachment conditions by approximately 10% (**Supplementary Figure 5E**). Regulation of expression of FOXM1 in cancer cells is diverse and can be both on the transcriptional as well as posttranscriptional level (Liao et al., 2018). Data from various solid tumors demonstrated that FOXM1 expression is substantially controlled by the PI3K-Akt pathway (Yao et al., 2018). Indeed, co-culture of BxPC-3 cells with platelets induced fast and strong phosphorylation of Akt at serine 473 (**Figure 5F**). Treatment with the Akt inhibitor MK-2206 reduced both FOXM1 expression as well as phosphorylation of Akt (**Figure 5G**), which was associated with attenuation of platelet-mediated anoikis resistance (**Figure 5H**).

Overall, these data demonstrated that platelets induced FOXM1 expression in detached cancer cells that was, at least partially, responsible for platelet-mediated anoikis resistance. Thus, targeting FOXM1 or inhibiting platelet-cancer cell interaction in circulating tumor cells might be a promising strategy to limit survival of CTCs and thereby reduce metastasis formation in pancreatic cancer.

## Discussion

Metastasis is the most lethal manifestation of cancer and most cancer patients die because of their metastatic disease (Chaffer and Weinberg, 2011). In particular, pancreatic cancer patients have a very poor overall survival rate after their disease has spread to distant organs (Siegel et al., 2024). A prerequisite for metastasis formation is detachment of tumor cells from the primary tumor, crossing of the endothelial cell barrier and survival in the blood stream despite immune cell attack (Gerstberger et al., 2023). Here, we provide evidence that platelets activate survival pathways within detached pancreatic cancer cells that are responsible for anoikis resistance. This is consistent with earlier reports showing that platelets closely interact with circulating tumor cells to promote their extravasation (Schumacher et al., 2013) or foster their survival in the peritoneal cavity (Haemmerle et al., 2017; Stone et al., 2012). Their interaction is reciprocal. Cancer cells are able to activate platelets via secretion of ADP (Haemmerle et al., 2016) that leads to platelet degranulation and secretion of pro-angiogenic, pro-tumorigenic and immunomodulatory factors (Klement et al., 2009; Maouia et al., 2020). In turn, this leads to cancer cell proliferation and facilitation of metastasis formation by inducing a mesenchymal-like phenotype in cancer cells (Labelle et al., 2011), by activating a YAP1-dependent transcriptional program (Haemmerle et al., 2017) or by guiding the formation of an early metastatic niche (Labelle et al., 2014). Interestingly, a recent study from the UK showed that regular aspirin use decreased the risk of PDAC, in particular in patients with diabetes mellitus (Buckland et al., 2024). Another retrospective study indicated that perioperative aspirin intake resulted in significantly better overall survival in pancreatic cancer patients due to an extended metastasis-free interval (Pretzsch et al., 2021). Platelets are likely to be the primary target of aspirin’s ability to inhibit platelet cyclooxygenase-1 (Lichtenberger and Vijayan, 2019). Despite the evidence supporting the role of platelets in tumor metastasis, prospective clinical trials have been scarce as anti-platelet medications may impair normal platelet function, resulting in bleeding complications. Here, we show that platelets interacting with detached cancer cells induce a FOXM1-dependent transcriptional program that enhanced cancer cell survival and promoted anoikis resistance. RNA sequencing analysis revealed a set of 24 genes consistently upregulated in tested pancreatic cancer cells by platelet interaction that were related to cell cycle progression. FOXM1 is a widely known transcription factor upregulated in a variety of human cancers (Kalathil et al., 2020; Myatt and Lam, 2007; Wierstra, 2013). In pancreatic cancer, FOXM1 is highest in the squamous subtype according to the dataset of Bailey et al. (Bailey et al., 2016) and high expression is associated with worse overall survival. Furthermore, it has been shown that miR-494 inhibits tumor growth and metastasis formation by targeting FOXM1-β-catenin signaling (Li et al., 2014). Another study highlighted the role of the FOXM1 isoform FOXM1c as a promotor of pancreatic cancer development and progression by enhancing uPAR signaling (Huang et al., 2014). Whether FOXM1 is associated with anoikis resistance in pancreatic cancer cells, has not been evaluated so far. Here, we show that FOXM1 mRNA and protein expression is significantly downregulated in detached pancreatic cancer cells and the extent of downregulation was inversely correlated with survival under low attachment conditions. Co-culture of platelets with detached pancreatic cancer cells significantly increased their survival capacity and upregulated FOXM1 expression as well as the expression of known FOXM1 target genes. In addition, inhibition of FOXM1 using the small molecule inhibitor FDI-6 (Gormally et al., 2014) reduced expression of FOXM1 transcriptional targets and diminished platelet-induced anoikis resistance. Similar results were obtained by siRNA-mediated FOXM1 downregulation. Lastly, our data suggested that FOXM1 expression is positively regulated by an activated Akt signaling pathway, which additionally contributed to platelet-mediated survival of pancreatic cancer cells. This is in line with a previous study that showed that FOXM1 is tightly regulated by the PI3K/Akt pathway in various cancer types (Yao et al., 2018). In conclusion, our in vitro experiments demonstrated that platelets upregulate FOXM1 expression in pancreatic cancer cells that contributes to platelet-mediated anoikis resistance. In the future, a detailed analysis is necessary to determine the primary signal provided by platelets that is responsible for the observed FOXM1 upregulation and to further investigate the biological function of FOXM1 in vivo. These analyses could, next to direct targeting of FOXM1, lead to innovative approaches to limit metastasis formation by disturbing the platelet-tumor cell interaction in pancreatic cancer patients, particularly in those with high platelet blood counts.

## Acknowledgments

We thank the core facility imaging (CFI) at the Institute of Molecular Medicine, in particular Nadine Bley and Danny Misiak, for excellent technical support with FACS, immunofluorescence analysis and NGS analysis. We thank Nadine Vollack-Hesse and Heidi Griesmann for their technical support regarding mouse handling, blood drawing and platelet isolation. This work was supported by research grants of the German Cancer Aid (70113190) to M.H. and the German Research Foundation (449501615, GRK 2751) to M.H. and T.G.

## Contributions

Conceptualization, A.E., T.G., M.H.; methodology, experiments, A.E., B.H., J.B., M.H.; data curation, A.E., M.H.; writing – original draft preparation, M.H.; writing – review and editing, A.E., T.G.; visualization, A.E., M.H.; project supervision, M.H.; funding acquisition, T.G., M.H.; All authors have read and agreed to the published version of the manuscript.

