## Supplementary Figures for "FOXM1 regulates platelet-induced anoikis resistance in pancreatic cancer cells"

### Supplementary information

**Supplementary figure 1: Anoikis rates change under low attachment versus attached conditions in pancreatic cancer cells. (A-E)** Pancreatic cancer cells were cultured under attached (AT) and low-attachment (LA) conditions for 72 hours and the % of dead (PI-positive) and living (PI-negative) CAPAN-1 (A), MIA Paca-2 (B), PA-TU-8988S (C), PA-TU-8988T (D) and AsPC-1 (E) cells were measured using flow cytometry (upper panel). In addition, increased apoptosis rates were confirmed using protein analysis and quantification of cleaved versus full length PARP (lower panel). GAPDH was used as a loading control. Bars and error bars represent mean values and the corresponding SEMs (n=3; ***$p < 0.001$).

**Supplementary figure 2: GSEA analysis and heat map of differentially regulated genes in Su.86.86 cells.** Significantly enriched pathways as evaluated by gene set enrichment analysis (GSEA) and heat map of overlapping genes between attached (AT) versus low attachment (LA) condition and LA with or without platelet (PLTS) co-incubation.

**Supplementary figure 3: FOXM1 expression in human pancreatic cancer. (A)** Analysis of the Bailey PDAC dataset (Bailey et al., Nature 2016) revealed a significant upregulation of FOXM1 expression in the squamous (S) compared to the pancreatic progenitor (PP), and ADEX (A) subtypes, whereas no difference could be observed compared to the immunogenic (I) subtype (*$p < 0.05$, **$p < 0.01$). **(B)** Survival analysis for PDAC patients with low FOXM1 (blue line, n = 49) versus high FOXM1 (red line, n = 47) expression (Bailey dataset, data accessed via https://r2.amc.nl, Log rank test).

**Supplementary figure 4: qRT-PCR validation of RNA sequencing results. (A-C)** mRNA expression of FOXM1-related genes after 48 hours of low attachment (LA) or attachment (AT) in Su.86.86 (A), IMIM-PC1 (B) and PA-TU-8988T (C) cells. (D) mRNA expression of FOXM1-related genes in IMIM-PC1 after 48h of low attachment with or without platelet (PLTS) co-incubation. RPLP0 was used as a reference gene. Bars and error bars represent mean values and the corresponding SEMs (n=3; ***$p < 0.001$, **$p<0.01$, *$p<0.05$, n.s. non-significant).

**Supplementary figure 5: FOXM1 expression in BxPC-3 cells. (A)** Number of PI-positive (dead) cells after low attachment BxPC-3 cultures in control conditions or with FDI-6 treatment. **(B)** Quantification of nuclear (NUC) and cytoplasmic (CYTO) FOXM1 expression in control cells, cells co-incubated with platelets, treated with FDI-6 or both (left); FOXM1 staining shown in B&W in indicated conditions (right). **(C)** FOXM1 expression in low attachment BxPC-3 cultures with or without platelet co-incubation and with or without siRNA transfection. **(D)** FOXM1 mRNA (upper panel) and protein expression (lower panel) in BxPC-3 cells with or without FOXM1 overexpression. RPLP0 was used as a reference gene and RPL7 was used as a loading control. **(E)** Percentage (%) of PI-positive (dead) cells after 72 hours of low attachment BxPC-3 cultures with or without FOXM1 overexpression. Bars and error bars represent mean values and the corresponding SEMs (n=3; ***$p < 0.001$, **$p<0.01$, *$p<0.05$, n.s. non-significant).

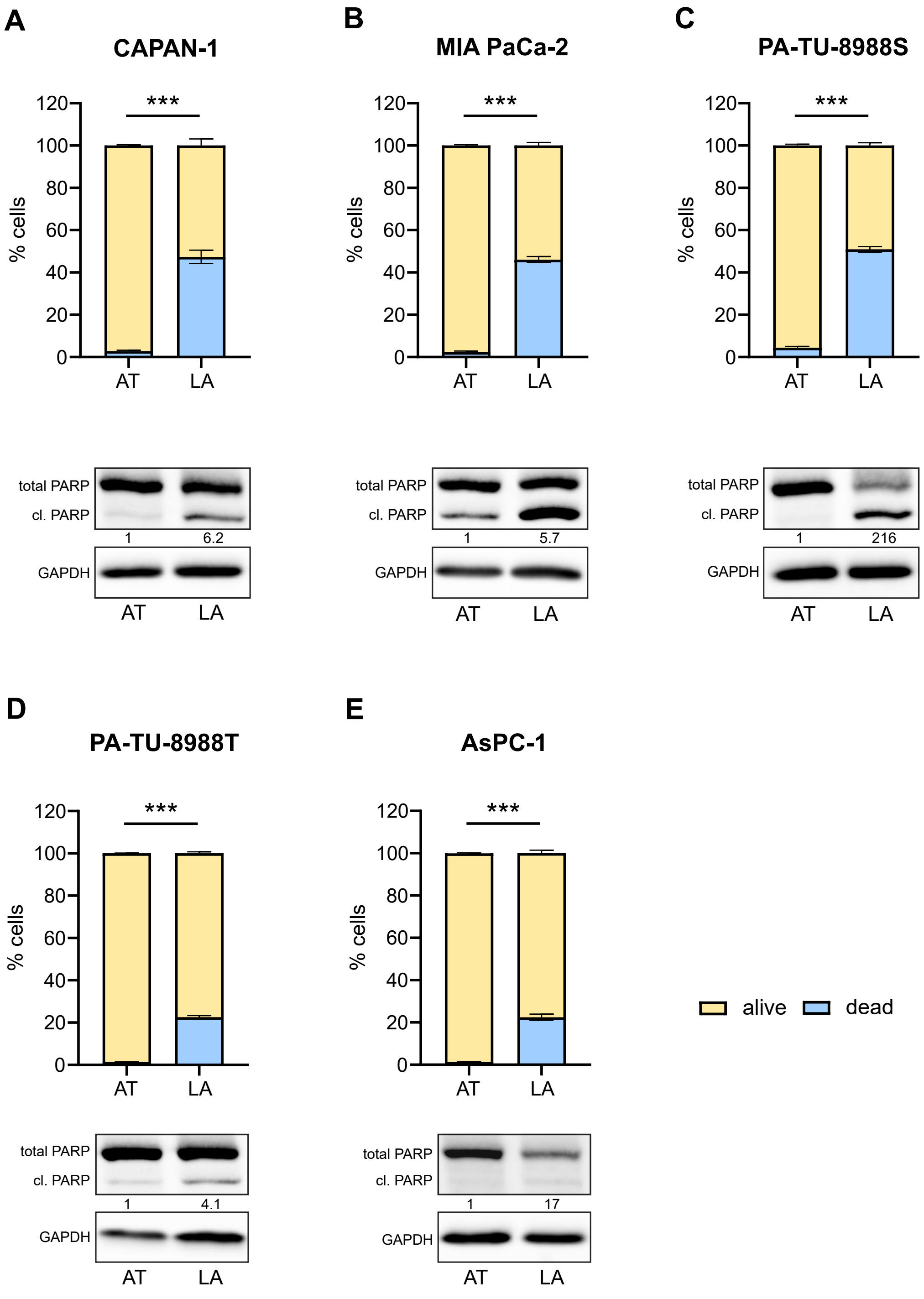


**Supplementary Figure 1**

A

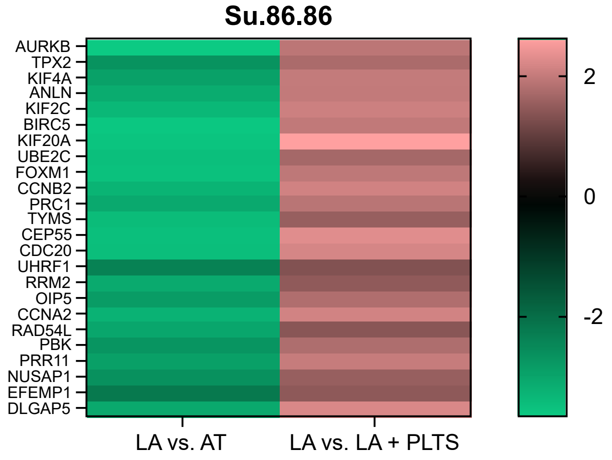


B

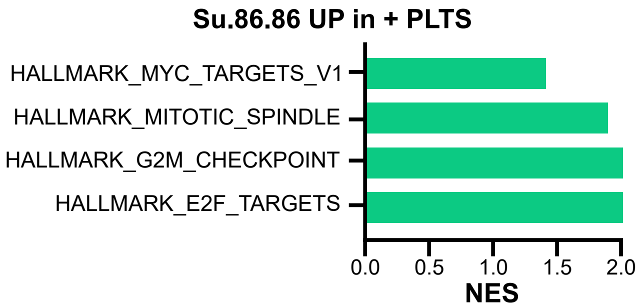


**Supplementary Figure 2**

A

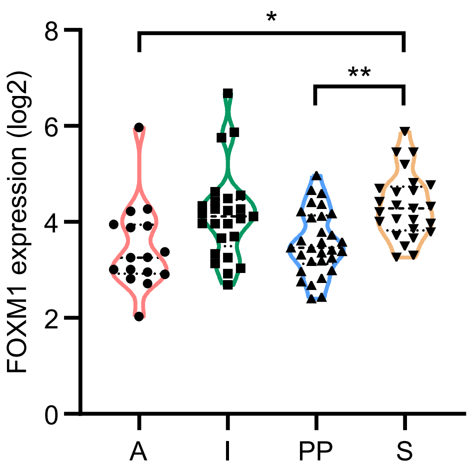


B

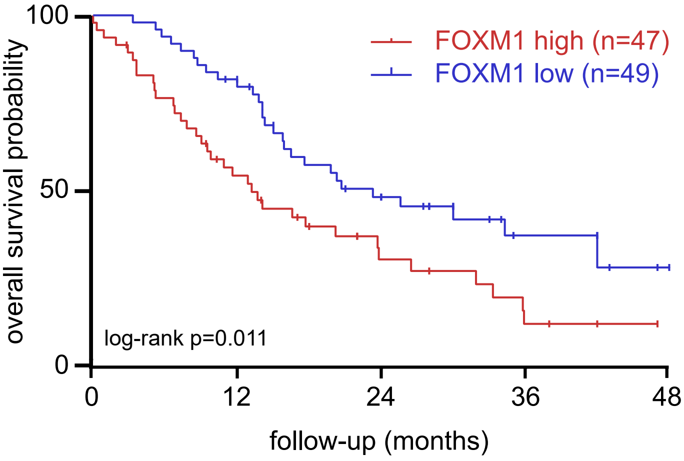


**Supplementary Figure 3**

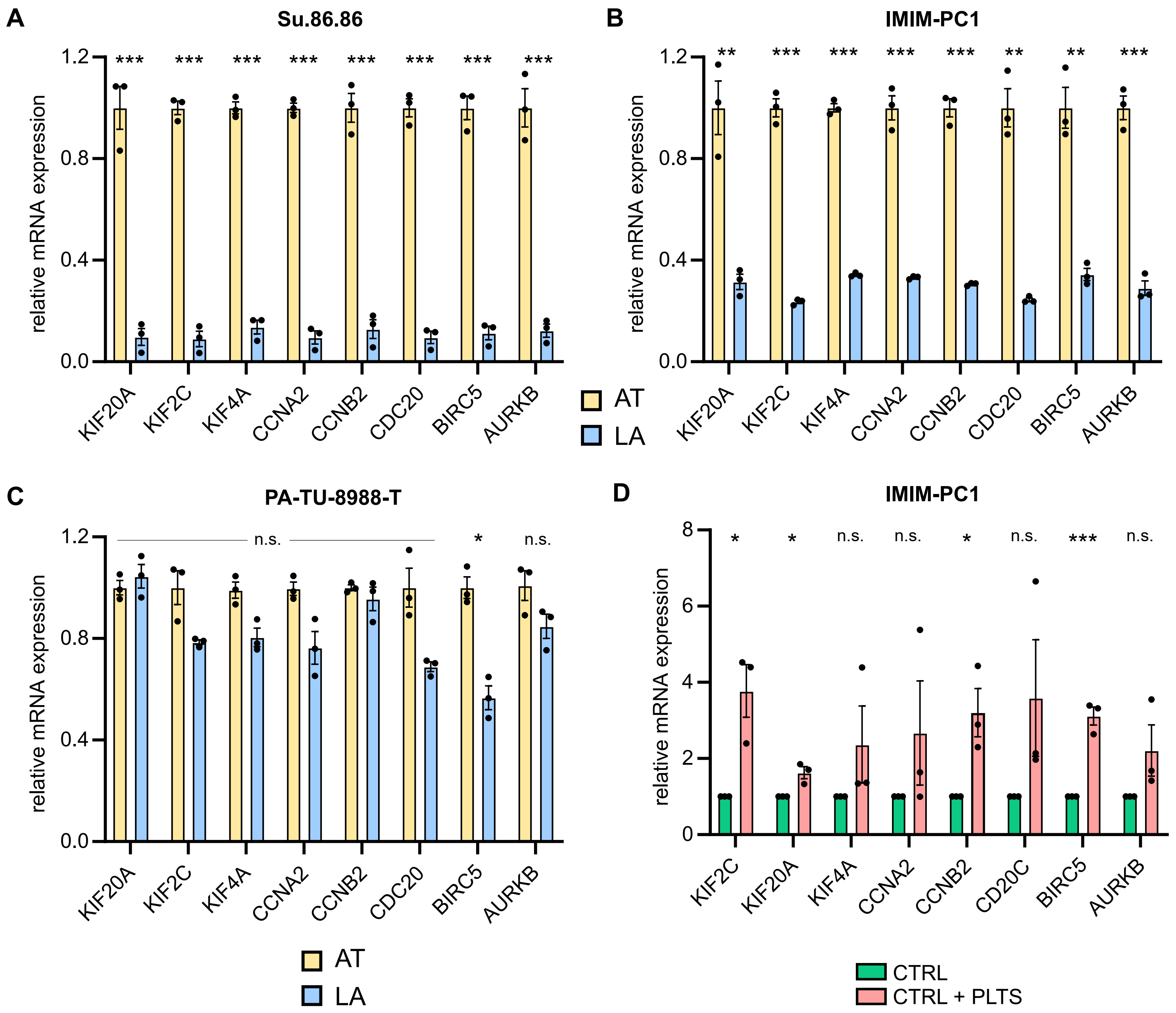


Supplementary Figure 4

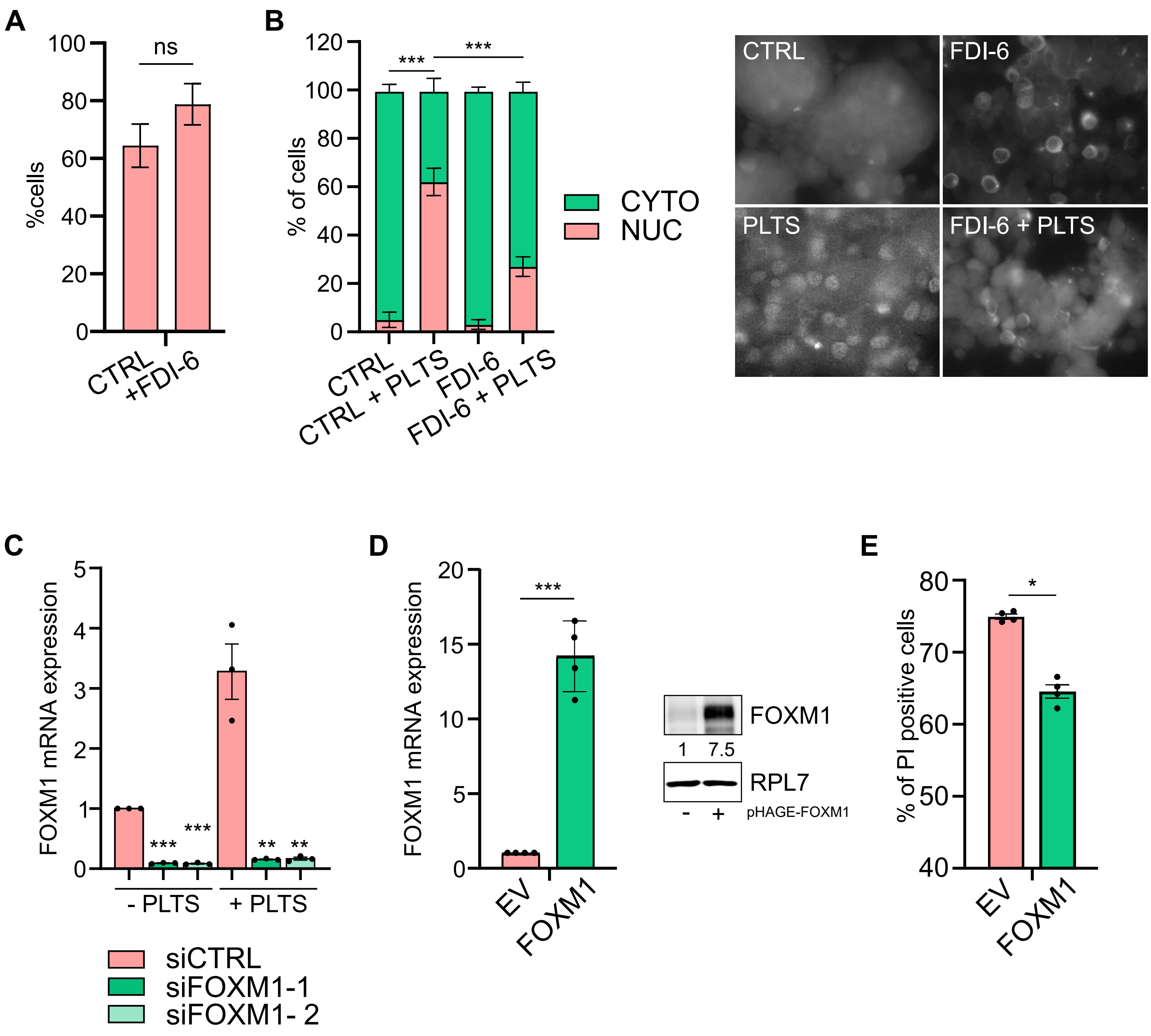


Supplementary Figure 5
